# Epidemiology of paediatric gastrointestinal colonisation by extended spectrum cephalosporin-resistant *Escherichia coli* and *Klebsiella pneumoniae* isolates in north-west Cambodia

**DOI:** 10.1101/173294

**Authors:** JJ van Aartsen, CE Moore, CM Parry, P Turner, N Phot, S Mao, K Suy, T Davies, A Giess, AE Sheppard, TEA Peto, NPJ Day, DW Crook, AS Walker, N Stoesser

**Affiliations:** Nuffield Department of Clinical Medicine and the National Institute for Health Research Oxford Biomedical Research Centre (NIHR-OxBRC), University of Oxford, Oxford, United Kingdom; Department of Clinical Infection, Microbiology and Immunology, Institute of Infection and Global Health, University of Liverpool, Liverpool, United Kingdom; Clinical Sciences, Liverpool School of Tropical Medicine, Liverpool, UK; Cambodia-Oxford Medical Research Unit, Angkor Hospitafor Children, Siem Reap, Cambodia; Centre for Tropical Medicine and Global Health, Nuffield Department of Medicine, University of Oxford, Oxford, UK; Angkor Hospital for Children, Siem Reap, Cambodia; Mahidol-Oxford Tropical Medicine Research Unit, Faculty of Tropical Medicine, Mahidol University, Bangkok, Thailand

**Author notes:** Corresponding author: Department of Microbiology/Infectious Diseases, John Radcliffe Hospital, Headley Way, Headington,OX3 9DU, United Kingdom.

## Abstract

Extended-spectrum cephalosporin resistance (ESC-R) in *Escherichia coli* and *Klebsiella pneumoniae* is a healthcare threat; high gastrointestinal carriage rates are reported from South-east Asia. Colonisation prevalence data in Cambodia are lacking. We determined gastrointestinal colonisation prevalence of ESC-resistant *E. coli* (ESC-R-EC) and *K. pneumoniae* (ESC-R-KP) in Cambodian children/adolescents and associated risk factors; characterised relevant resistance genes, their genetic contexts, and the genetic relatedness of ESC-R strains using whole genome sequencing (WGS). Faeces and questionnaire data were obtained from individuals <16 years in northwestern Cambodia, 2012. WGS of cultured ESC-R-EC/KP was performed (Illumina). Maximum likelihood phylogenies were used to characterise relatedness of isolates; ESC-R-associated resistance genes and their genetic contexts were identified from *de novo* assemblies using BLASTn and automated/manual annotation. 82/148 (55%) of children/adolescents were ESC-R-EC/KP colonised; 12/148 (8%) were co-colonised with both species. Independent risk factors for colonisation were hospitalisation (OR: 3.12, 95%, CI [1.52-6.38]) and intestinal parasites (OR: 3.11 [1.29-7.51]); school attendance conferred decreased risk (OR: 0.44 [0.21-0.92]. ESC-R strains were diverse; the commonest ESC-R mechanisms were *bla*_CTX-M_ 1 and 9 sub-family variants. Structures flanking these genes were highly variable, and for *bla*_CTX-M-15,-55 and-27_, frequently involved IS*26*. Chromosomal *bla*_CTX-M_ integration was common in *E. coli*. Gastrointestinal ESC-R-EC/KP colonisation is widespread in Cambodian children/adolescents; hospital admission and intestinal parasites are independent risk factors. The genetic contexts of *bla*_CTX-M_ are highly mosaic, consistent with rapid horizontal exchange. Chromosomal integration of *bla*_CTX-M_ may result in stable propagation in these community-associated pathogens.

## MAIN TEXT

### INTRODUCTION

*Escherichia coli* and *Klebsiella pneumoniae* are two bacterial pathogens of the Enterobacteriaceae family that can cause a wide spectrum of clinical disease, ranging from cystitis and intra-abdominal abscesses to sepsis. Both species also asymptomatically colonise the gastrointestinal tract, a reservoir that assists in the acquisition and spread of antimicrobial resistance (AMR)(1, 2). The increasing prevalence of AMR worldwide is reducing the efficacy of our limited armamentarium of empirical broad-spectrum antibiotics, such as extended-spectrum cephalosporins (ESCs), resulting in increased healthcare costs and mortality(3-5).

Recent reports from South-east Asia show substantial variation between country and cohort in gastrointestinal colonisation by Enterobacteriaceae possessing Ambler class A extended spectrum beta-lactamases (ESBLs) and/or class C AmpC enzymes, which can hydrolyse third and fourth generation cephalosporins. In the Lao People‘s Democratic Republic, for example, 23% of pre-school children carried these strains, in contrast to a much higher prevalence of 65.7% in a rural Thai adult population(6-8). Data describing the prevalence and mechanisms of antibiotic resistance in Cambodia are limited to only a few studies. Vlieghe and colleagues found 49.7% of Enterobacteriaceae from blood cultures in Phnom Penh from 2007-2010 were cefotaxime-resistant, mostly due to CTX-M-15 and CTX-M-14 enzymes(9). Studies from 2004/5 and 2007-2011 identified ESC resistance in 36-44% of urinary tract infection isolates(10, 11). The gastrointestinal colonisation prevalence of ESC-resistant (ESC-R) *E. coli* and *K. pneumoniae* in Cambodia has previously only been investigated in hospitalised neonates(12).

This study aimed to: (i) estimate the prevalence of gastrointestinal colonisation with ESC-resistant *E. coli* (ESC-R-EC) and *K. pneumoniae* (ESC-R-KP) in Cambodian children and adolescents, and the molecular mechanisms responsible; (ii) investigate risk factors for ESC-R colonisation; (iii) determine genetic relatedness of ESC-R strains.

## RESULTS

### Sampling, culture and basic demographics

In total, 196 faecal samples were obtained from a consecutive subset of children/adolescents enrolled in a helminth prevalence study. 48 samples were excluded from this study because of: (i) lack of specific consent for wider use of the faecal samples beyond the helminth survey (n=36); (ii) no epidemiological data records (n=1); (iii) no (n=3) or poor (n=5) growth on culture; or (iv) replicate samples for the same patient (n=3), leaving 148 samples/individuals for analysis.

Overall, 184 distinct colony types grew within the cefpodoxime inhibition zones; 141 were pink (presumed *E. coli*) and 43 were blue (presumed *Klebsiella* spp., *Enterobacter* spp. or *Citrobacter* spp.). All pink colonies but only 22/43 (54%) blue colonies were confirmed as phenotypically ESC-R using BSAC methods. All 163 confirmed ESC-R isolates were sequenced; two failed and were excluded from further analysis. Of the 161 sequences, *in silico* species identification confirmed 135 (84%) isolates were *E. coli*, 18 (11%) *K. pneumoniae*, and 8 (5%) *Enterobacter* spp. 38 *E. coli* isolates and one *K. pneumoniae* isolate were genetically sufficiently closely related to another isolate obtained from the same patient sample to be considered as the same strain (defined as ≤5 chromosomal SNVs); these were also excluded leaving 122 isolates for analysis. None of the 148 faecal samples yielded imipenem resistant colonies.

Participants were median 4.2 years old (interquartile range: 1.1-8.8) at sample collection; 70/148 (47%) were male. 70/147 (48%; 1 missing) were inpatients at sample collection. Although most were from Siem Reap province (99/148 [67%]), the hospital catchment is such that the remainder were recruited from 10 other provinces. 16/148 (11%) were clinically malnourished, and 23/148 (16%) had ≥1 underlying chronic medical condition including HIV (n=5), haematological disease (n=3), congenital cardiac disease (n=5), tuberculosis (n=4), and asthma (n=2)(Table 1).

**Table 1.**
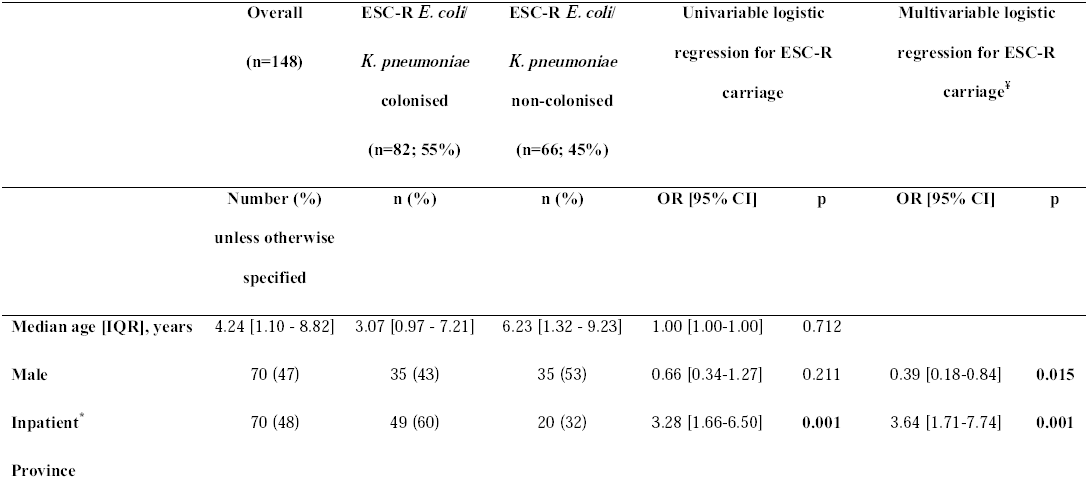
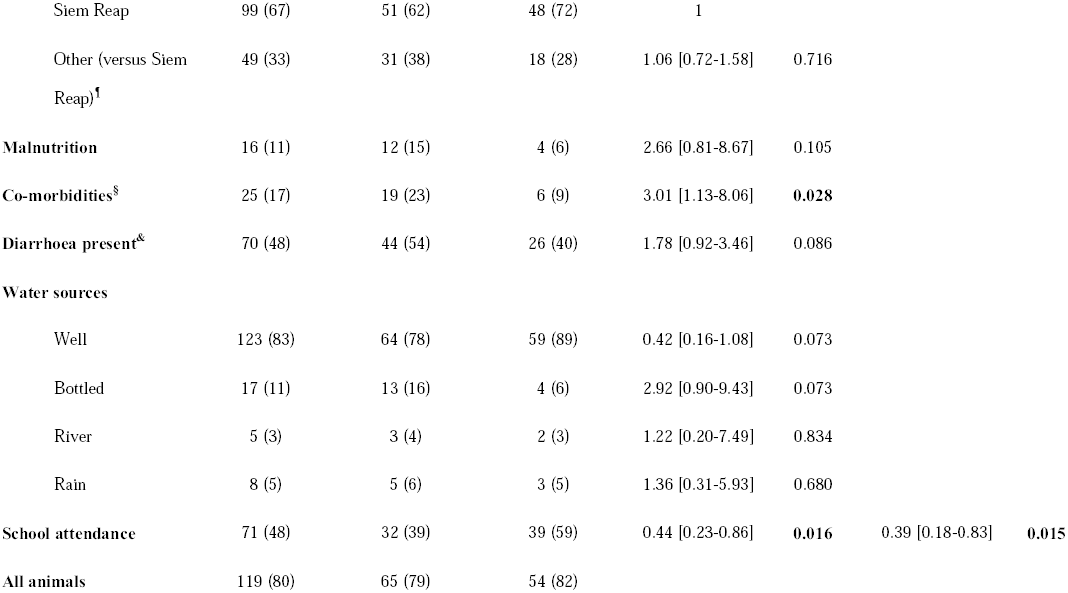
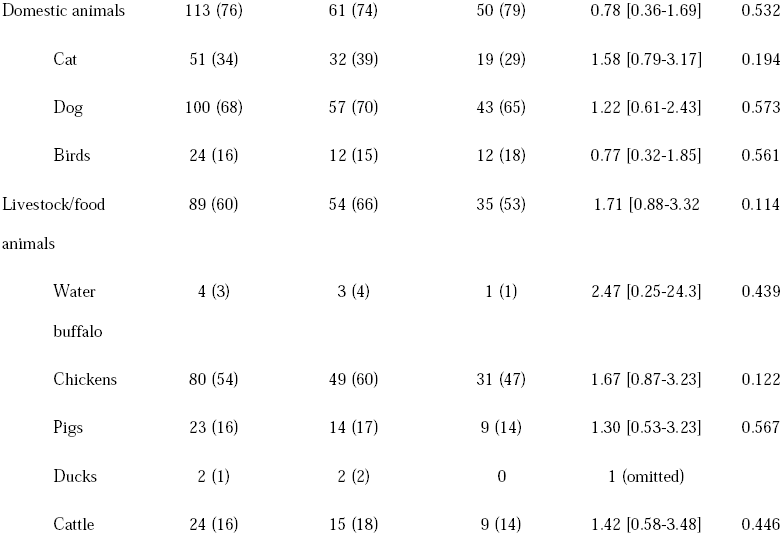
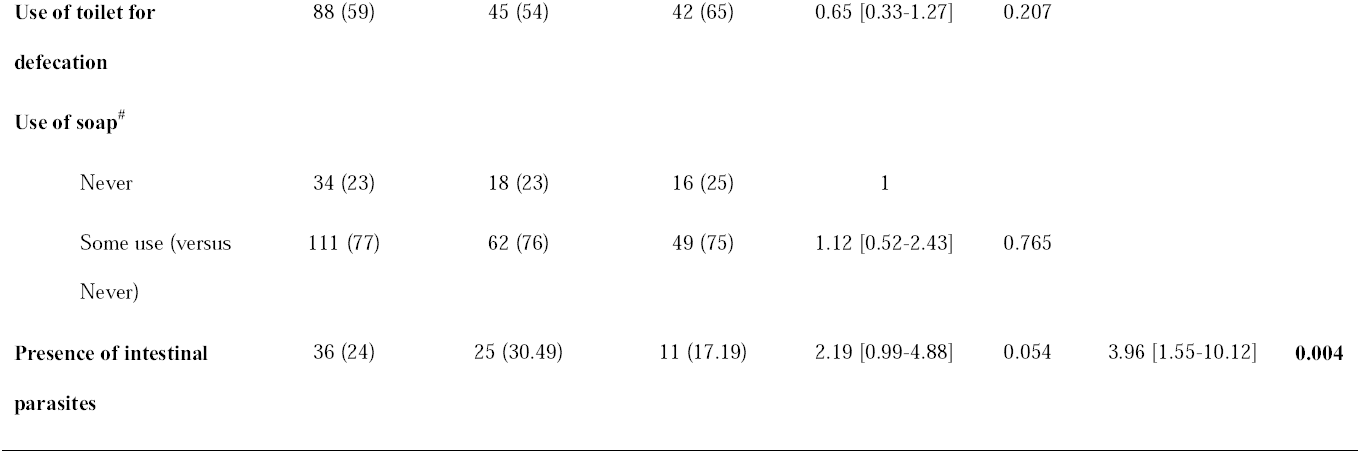
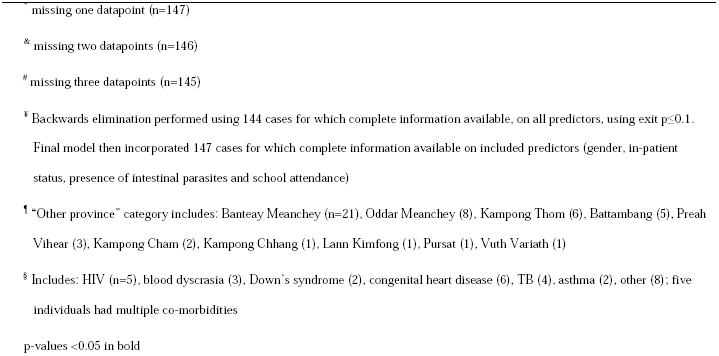
Clinical and epidemiological details of all 148 participants, also categorised by presence/absence of gastrointestinal colonisation with ESC-resistant *E. coli* and/or *K. pneumoniae*, and multivariable logistic regression outcomes.

### Prevalence of and risk factors for colonisation with ESC-R EC and/or ESC-R-KP

A total of 114 confirmed ESC-R-EC (n=97) and ESC-R-KP (n=17) remained in the analysis and were carried by 82/148 participants, giving a combined ESC-R-EC/KP prevalence of 55% (95% CI: 47%-64%); 53% for ESC-R EC (79/148 patients; 95% CI: 45%-62%) and 10% for ESC-R KP (15/148 patients; 95% CI: 6%-16%). Co-colonisation with both ESC-R-EC and ESC-R-KP was observed in 12/82 (15%). Independent risk factors for ESC-R-EC/KP colonisation included being a current inpatient (OR=3.64; 95% CI [1.71-7.74), p=0.001) and the presence of faecal parasites (OR=3.96 [1.55-10.13], p=0.004). ESC-R-EC/KP colonisation was lower in males (OR=0.39 [0.18-0.84], p=0.015) and in those attending school (OR=0.39 [0.18-0.83], p=0.015)(Table 1).

### Sequence type, Ambler class and genetic mechanisms of ESC-R

The 97 ESC-R-EC isolates came from 33 known and 6 novel STs (Fig.1, for details see Table S1). 22% (17/79) of patients were colonised by at least two different ESC-R-EC STs, although this may underestimate diversity as only a small number of colonies (≤3) were sampled per patient(25). The 17 ESC-R-KP strains came from 11 known and 3 novel STs (n=4 isolates) (Fig.2, Table S2). Two patients were colonised by two different ESC-R *K. pneumoniae* STs (2/15, 13%).

**Figure 1.**
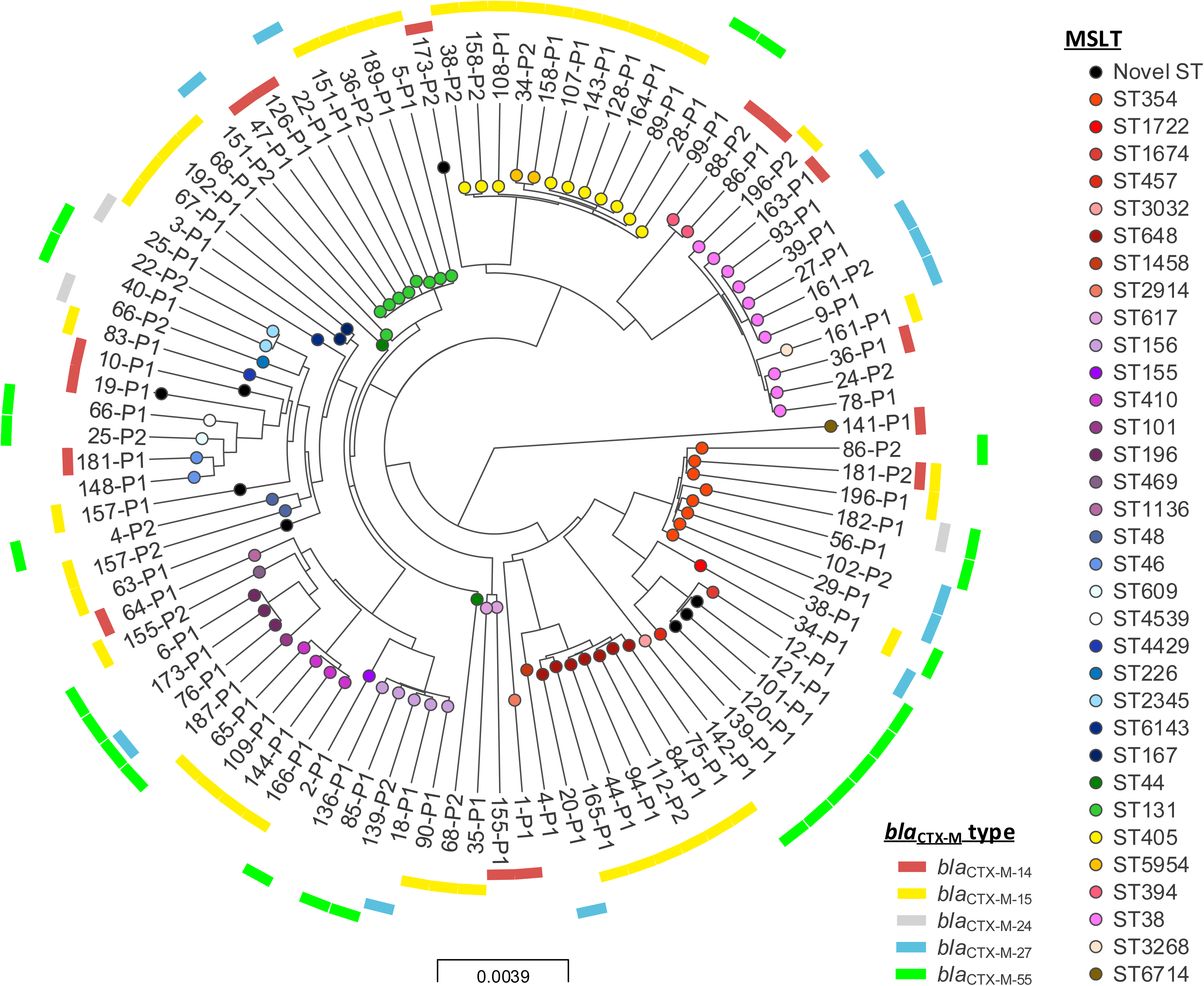
Phylogeny of study *Escherichia coli* isolates. Interactive map of geographic locations and genetic attributes can be visualised at: https://microreact.org/project/By8bf5ajg

**Figure 2.**
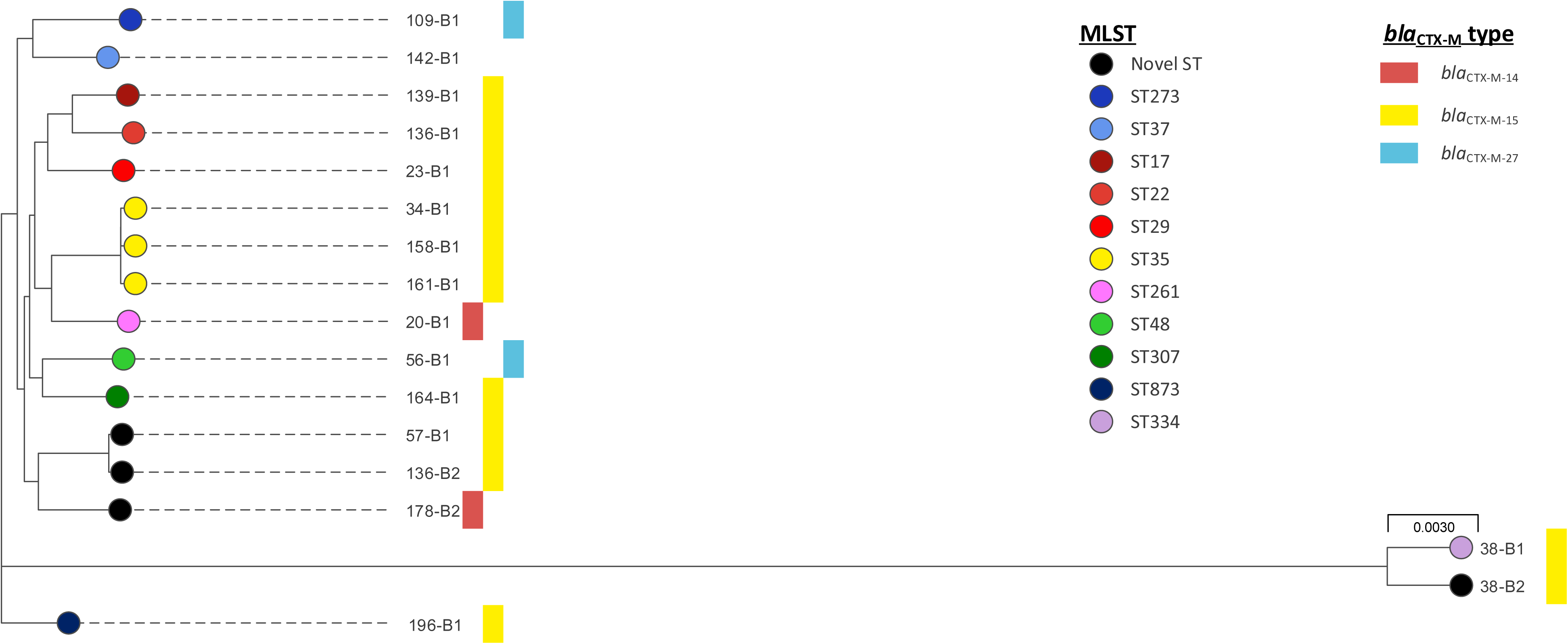
Phylogeny of study *Klebsiella pneumoniae* isolates. Interactive map of geographic locations and genetic attributes can be visualised at: https://microreact.org/project/Hy_yQcaog

In total, 77% (88/114) and 23% (26/114) of isolates displayed Ambler class A or C phenotypes, respectively. Neither species were associated with Ambler class A (76% [74/97] versus 82% [14/17]) or class C (23% [23/97] versus 18% [3/17]; Fishers exact test; p=0.759). In all class A isolates the phenotype could be explained by the presence of one (84/88, 95%) or two (4/88, 5%) *bla*_CTX-M_ genes; *bla*_SHV_ (12/88, 12%) and *bla*_VEB_ (1/88, 1%) occurred less commonly. Class C gene families were only identified in 39% (10/26) of phenotypically class C isolates: specifically *bla*_CMY-2_ (8/26, 31%) or *bla*_DHA_ (2/26, 8%). In the remaining 16 isolates, the genetic basis for the class C phenotype was unclear; of note, however, *ampC* promoter mutations were not assessed. 111 *bla*_CTX-M_ genes were found in 94% (107/114) of ESC-R-EC/KP, with two separate alleles identified in 4% of isolates (4/114). The most frequently identified allele was *bla*_CTX-M-15_ (53/111, 48%), followed by: *bla*_CTX-M-55_ (24/111, 22%), bla_CTX-M-14_ (17/111,15%), *bla*_CTX-M-27_ (14/111, 13%) and *bla*_CTX-M-24_ (3/111, 3%). Two different *bla*_CTX-M_ alleles were found in 21% (18/82) of individuals carrying ESC-R-EC/KP. *bla*_SHV_ genes were identified in 15/17 *K. pneumoniae*, including *bla*_SHV-1/SHV-1-like_ (3/15, 20%), bla_SHV-11/SHV-11-like_ (4/15, 27%)*, bla*_SHV-27-like_ (1/15, 7%)*, bla*_SHV-28_ (1/15, 7%)*, bla*_SHV-33_ (3/15, 20.0%) and *bla*_SHV-83_ (1/15, 7%), *bla*_SHV-99-like_ (1/15, 7%) and *bla*_SHV-142_ (1/15, 7%). All eight *bla*_CMY-2/CMY-2-like_ genes were found in *E. coli*. The study population carriage prevalence of common genetic mechanisms encoded by ESC-R EC/KP was therefore: 53% *bla*_CTX-M_ (78/148), 9% *bla*_SHV_ (14/148), 1% *bla*_VEB_ (1/148), 5% *bla*_CMY-2_ (8/148), 1% *bla*_DHA_ (2/148). Two individuals (1%) carried isolates with *bla*_OXA-48_ (one *K. pneumoniae* ST48 and one *E. coli* ST648); no other carbapenem resistance mechanisms were identified.

### Genetic context of ESC-R genes

For the 41 *E. coli* harbouring *bla*_CTX-M-15_, it was chromosomally located in five cases (12%), and likely in plasmid contexts in two; in the remaining cases it was not possible to determine wider chromosomal/plasmid location (Table 2). One isolate (38P1) harboured short contigs containing truncated *bla*_CTX-M-15_, leaving 40 cases in which to evaluate the immediate flanking contexts surrounding the *bla*_CTX-M_ gene. All contained IS*Ecp1* upstream of *bla*_CTX-M-_ 15, but with considerable evidence of additional mobilisation events/mosaicism (Table 2). In particular, IS*Ecp1* was truncated by IS*26* at 24, 497, 524, 1067, 1173, 1421, or 1489bp in 13 isolates, consistent with at least seven IS*26*-associated insertion events within IS*Ecp1*(Fig.3). Another 13 IS*Ecp1* elements were truncated by contig breaks, without any specific associated genetic signatures, although contig breaks are frequently due to repeat structures and may therefore have represented additional disruption events. One isolate had an intact IS*Ecp1* element, without any wider flanking upstream context. The 13 cases with an intact IS*Ecp1* were consistently flanked by variable lengths of Tn*2*, which was truncated by an IS*26* right IRR in 2/7 evaluable cases (and by an unknown sequence in the other 5/7). Two isolates had a complete Tn*2* structure interrupted by IS*Ecp1*-*bla*_CTX-M-15_ (TCTCA-TCTCA and TTTTA-TAAAA target site sequences [TSSs] respectively)(Fig.3). Overall, genetic contexts of *bla*_CTX-M-15_ were consistent with integration and mobilisation of IS*Ecp1*-*bla*_CTX-M-15_ within a Tn*2* element, as previously described(26), with subsequent rearrangement events facilitated by IS*26* and perhaps other ISs(27)(Table S3).

**Figure 3.**
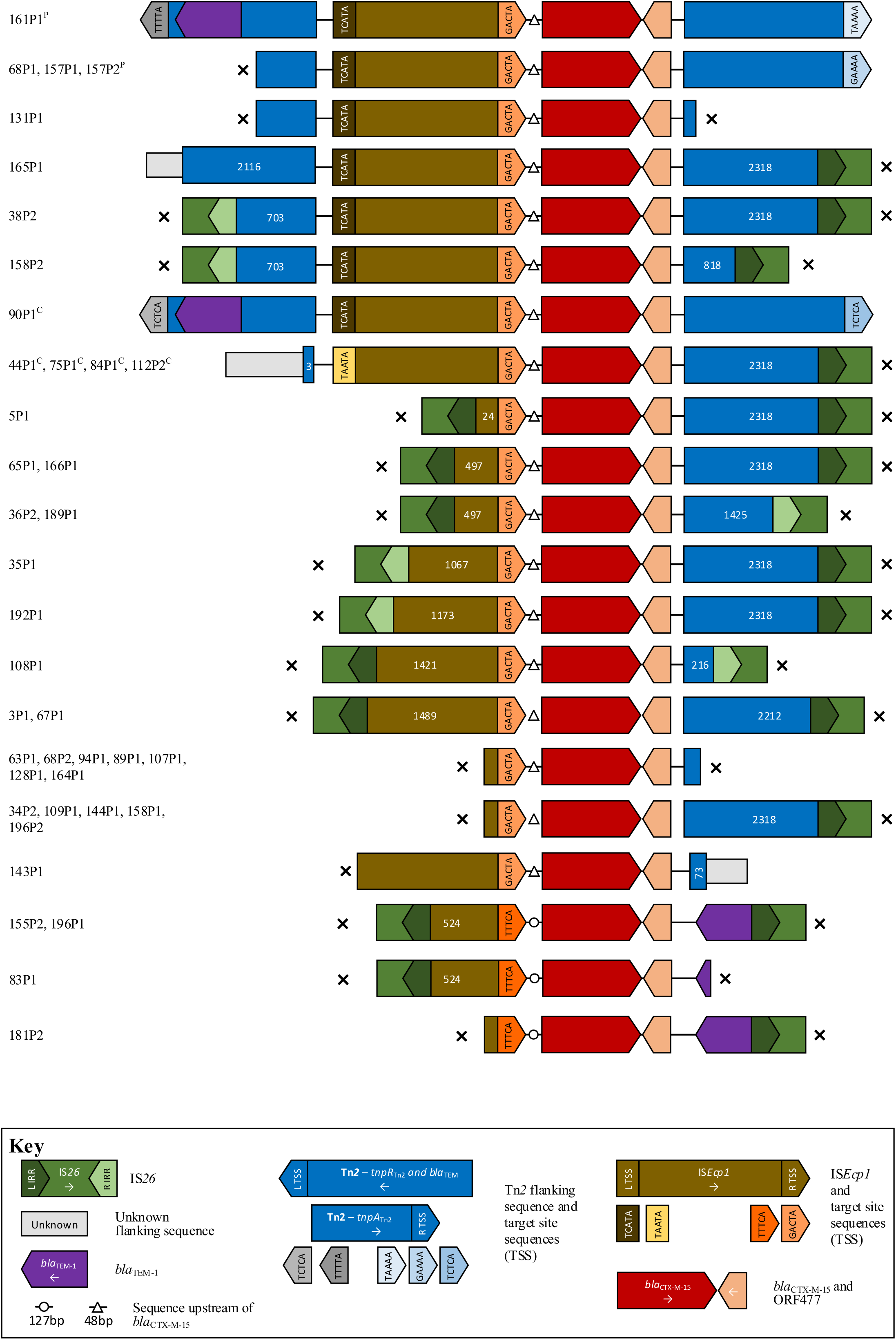
Schematic of aligned genetic contexts for *bla*_CTX-M-15_ in study *Escherichia coli*. Features of interest are highlighted in the figure key. White numbers within open reading frames denote truncated sequence length (bp). Isolates harbouring this genetic context are listed to the left of the figure. “x” denotes contig breaks. ^P^ denotes plasmid contexts; ^c^ chromosomal contexts.

**Table 2.**
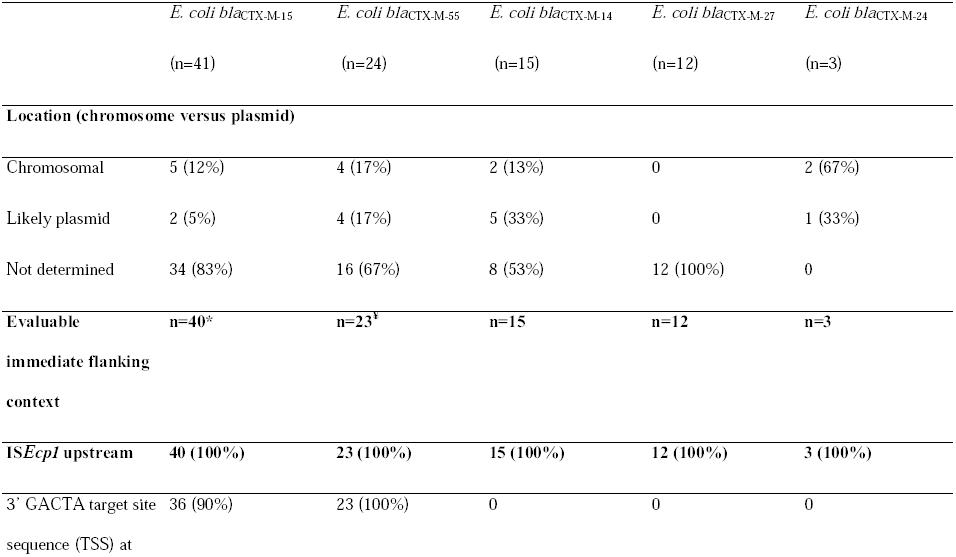
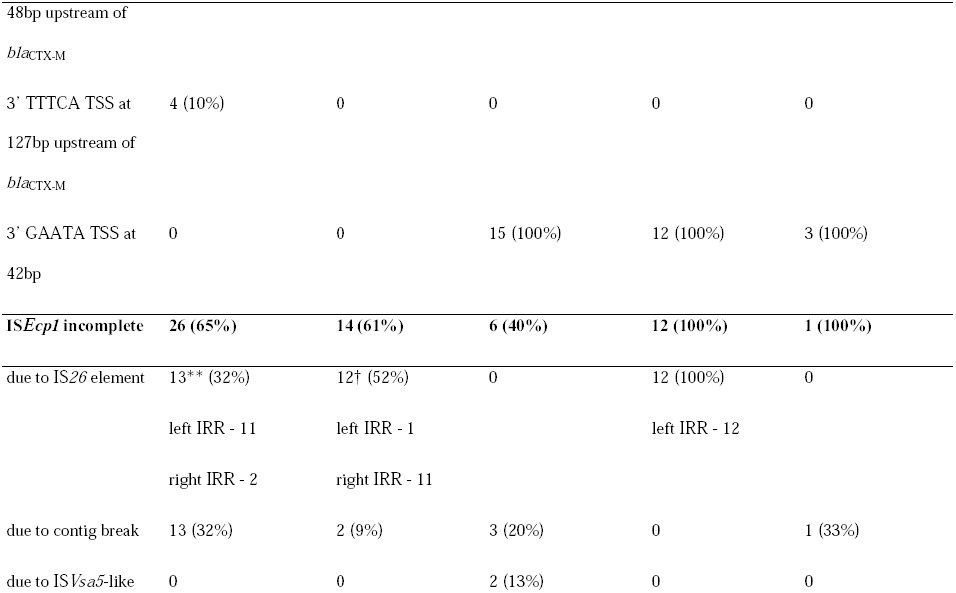
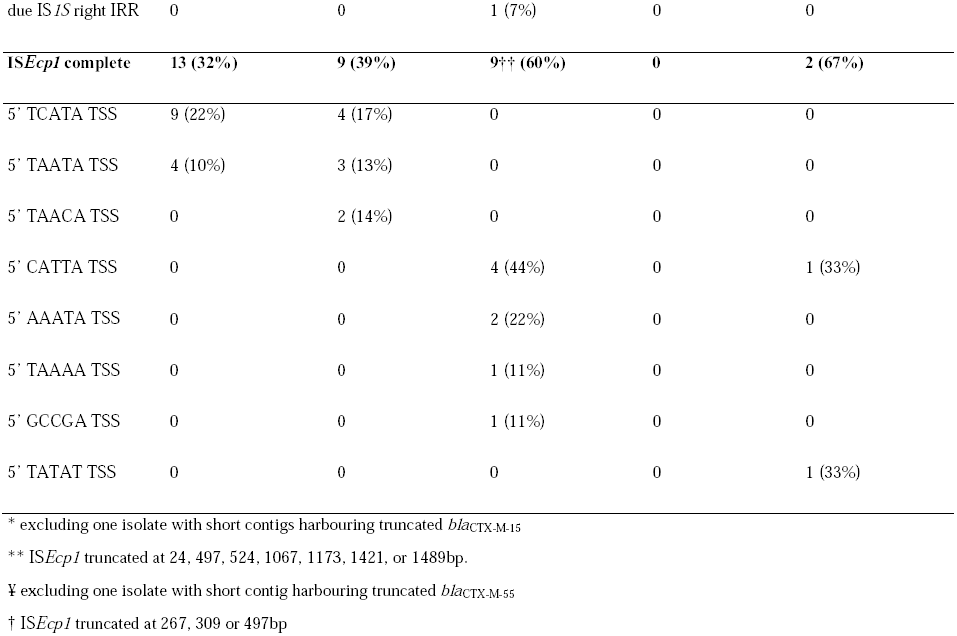
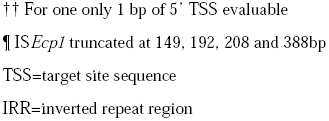
Summary of genetic co 419 ntexts of *bla*_CTX-M_ in *E. coli*.

For the 24 *E. coli* harbouring *bla*_CTX-M-55_, it was chromosomally located in 4 (17%), plasmid in 3 (13%) and unknown in 16 (67%). One contig contained a truncated *bla*_CTX-M-55_, leaving 23 evaluable contexts. Similar to *bla*_CTX-M-15_, it was invariably associated with IS*Ecp1* upstream of *bla*_CTX-M-55_ (Fig.4), which was often incomplete, representing at least 3 different IS*26*-associated IS*Ecp1* disruption events (Table 2). Intact IS*Ecp1* were flanked by variable lengths of Tn*2* sequence, apart from 120P1 where the contig was truncated immediately at the 5‘ end of IS*Ecp1*. One isolate (2P1) had the same *bla*_CTX-M_/Tn*2* unit as for *bla*_CTX-M-15_ (but with TACTC-TAAAA), consistent with the evolution of *bla*_CTX-M-55_ from *bla*_CTX-M-15_ (1 SNV difference) within this unit (Figs.3, 4).

**Figure 4.**
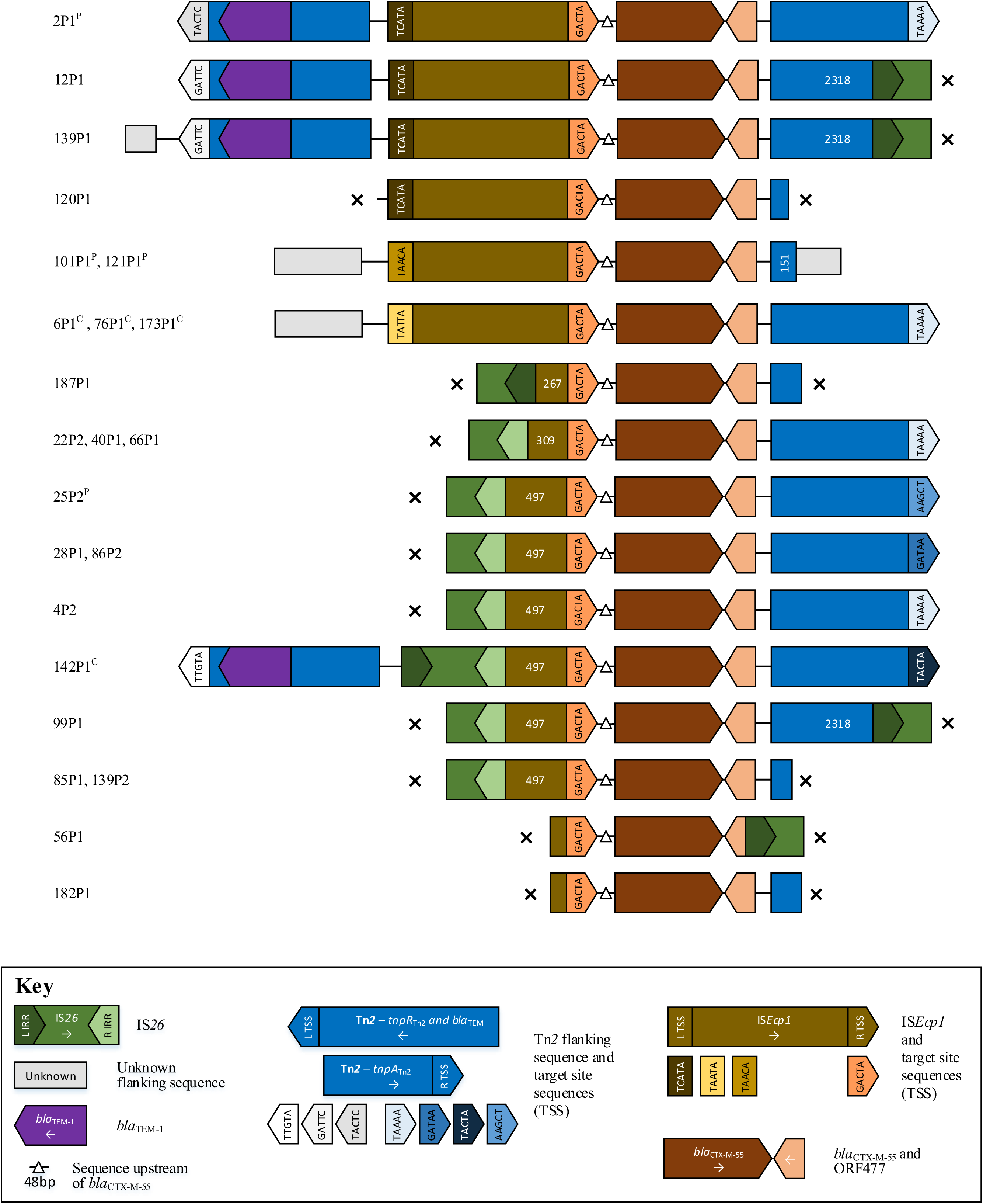
Schematic of aligned genetic contexts for *bla*_CTX-M-55_ in study *Escherichia coli*. Features of interest are highlighted in the figure key. White numbers within open reading frames denote truncated sequence length (bp). Isolates harbouring this genetic context are listed to the left of the figure. “x” denotes contig breaks. ^P^ denotes plasmid contexts; ^c^ chromosomal contexts.

For the 15 *E. coli* harbouring *bla*_CTX-M-14_, it was chromosomally located in 2 (13%) cases, plasmid-associated in 5 (33%), and unknown in 8 (53%). Again, it was invariably associated with IS*Ecp1*, but more often complete and with different mechanisms of disruption (2 IS*Vsa5-like* sequence, one IS*1S* R IRR). All cases had an IS*903* element at the 3‘ end of *bla*_CTX-M-14_; this had been disrupted in 6 cases, with additional contig breaks in 5 cases (Fig.5). Two of three *E. coli bla*_CTX-M-24_ contexts were chromosomal, with flanking contexts similar to *bla*_CTX-M-14_ (Fig.S1). In the 12 *bla*_CTX-M-27_ cases, the *ISEcp1* element had been disrupted by an IS*26* L IRR in all contexts, at 149, 192, 208 and 388bp, but the wider genetic context of this structure was indeterminable in all cases (Fig.S2).

**Figure 5.**
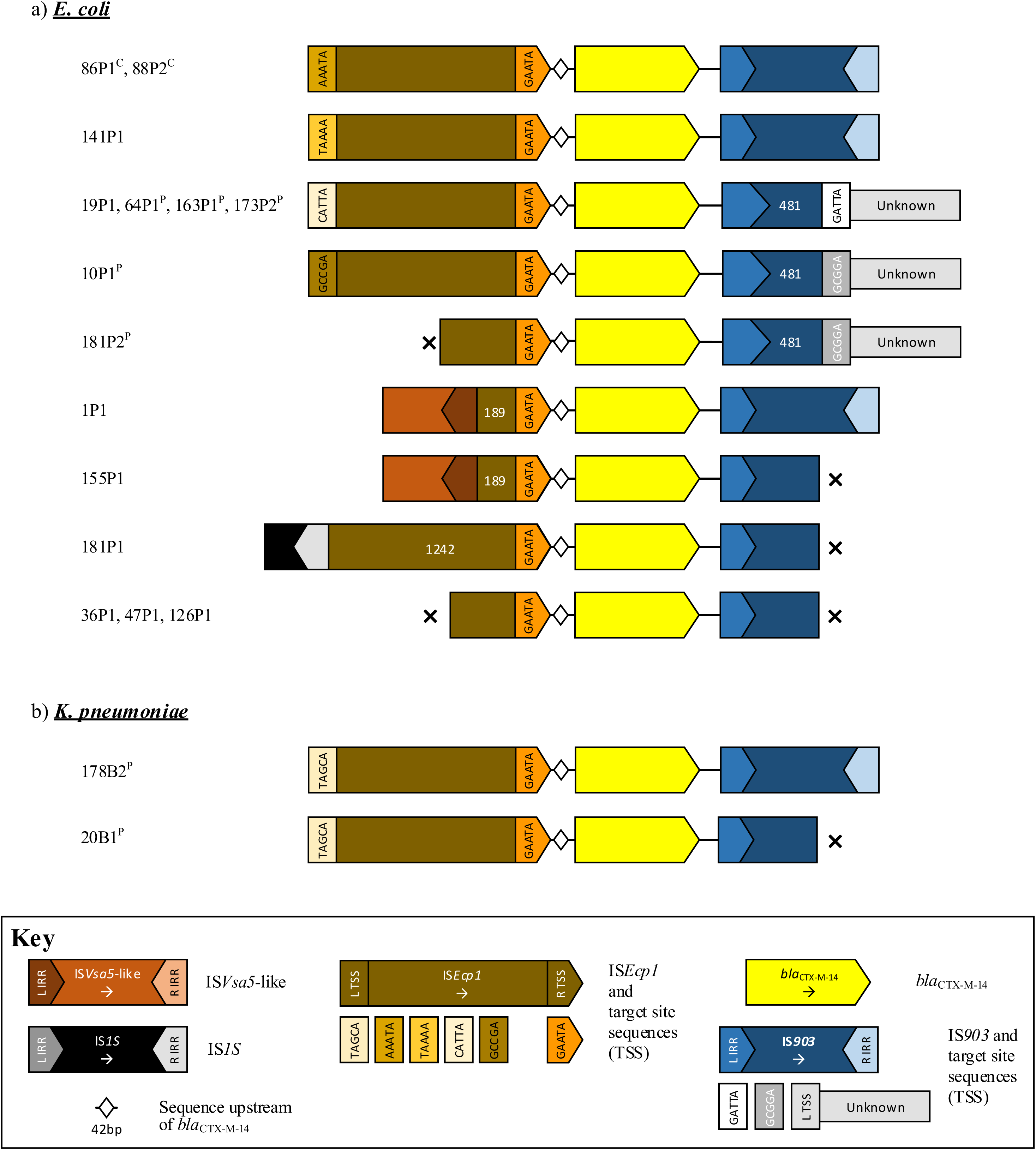
Schematic of aligned genetic contexts for *bla*_CTX-M-14_ in study *Escherichia coli* and *Klebsiella pneumoniae*. Features of interest are highlighted in the figure key. White numbers within open reading frames denote truncated sequence length (bp). Isolates harbouring this genetic context are listed to the left of the figure. “x” denotes contig breaks. ^P^ denotes plasmid contexts; ^c^ chromosomal contexts.

Overall, *bla*_CTX-M_ was chromosomal in 13/92 cases (14%; 13/25 [52%] cases where plasmid versus chromosomal location could be assessed), suggesting that CTX-M genes may be incorporated chromosomally and indiscriminately in significant numbers of colonising *E. coli*, with possible implications for their stable propagation within the wider *E. coli* population.

For *K. pneumoniae*, 12 isolates harboured *bla*_CTX-M-15_, in a plasmid-associated context in 9/12 cases, and an unknown context in 3/12 cases. Three isolates harboured a complete *bla*_CTX-M-15_ /Tn*2* complex with GTTAA-GTTAA TSS, most consistent with a direct transposition of this element into a plasmid context. In the other isolates, the IS*Ecp1*-*bla*_CTX-M-15_-ORF477 was flanked by variable stretches of Tn*2*-associated sequence identical to that found in the *E. coli* isolates, and similarly truncated either as a result of contig breaks, or by IS*26* inverted repeats, consistent with between species and within species mobilisation (Fig.S3).

Four *K. pneumoniae* isolates harboured *bla*_CTX-M-9_ group genes; two of these (*bla*_CTX-M-14_) shared the same IS*Ecp1* (Fig.5) and ∼18kb upstream flanking plasmid sequence; and two (*bla*_CTX-M-27_) an IS*Ecp1* element truncated at position 1499 by an IS*26* L IRR (Fig.S2).

## DISCUSSION

We observed significant gastrointestinal carriage prevalence of both ESC-R-EC and ESC-R-KP in Cambodian children sampled in 2012; approximately one in twelve children was co-colonised with ESC-R strains of both species. A wide diversity of ESC-R strain types was observed, including several genotypes categorised as “high risk” clones, such as *E. coli* STs 38, 405, 131, 354 and 648(13). The predominant ESC-R genotypic mechanism was *bla*_CTX-M_, with the major allelic variants being those widely described elsewhere in Asia (Group 1: *bla*_CTX-M-15,-55_, Group 9: *bla*_CTX-M-14,-24,-27_). Approximately one-third of the Cambodian population is <18 years old, so this group may be acting as a significant reservoir for the spread of anti-microbial resistant organisms. We did not observe particularly high rates of colonisation with carbapenem-resistant isolates (2 (1%) individuals), but one of these was an out-patient, with an OXA-48 *E. coli* isolate, and without any known chronic health problems, suggesting that there may be some carriage of carbapenem-resistant isolates in the community. Further assessment of the extent of carriage of carbapenem-resistant EC/KP in this context is warranted.

Independent risk factors for colonisation included inpatient status, consistent with transmission within hospital, and/or selection of these organisms from low-level carriage by the use of antibiotics on admission given the high burden of infectious diseases in this region. Infection control (IC) in resource-limited settings remains challenging, and despite improvements within the study hospital(14), recent longitudinal surveillance within the neonatal care unit identified high rates of import of ESC-R-KP (62% colonised on admission) as well as nosocomial acquisition (23%)(12). In-patient acquisition of ESC-R-EC/KP has also been identified as a major problem in other low/middle-income settings(15). The specific effect of faecal parasites on gut microbiota is not well-studied, but they are thought to significantly perturb microbial diversity(16). Helminth infestation may also result in inappropriate antimicrobial use, including antibiotics, perhaps leading to secondary colonization with drug-resistant commensals. The decreased risk associated with school attendance has been observed in a previous study in Spain(17), and may represent a proxy marker for increased socio-economic status, and parental levels of education, which were not evaluated here, but may translate into better awareness of appropriate antibiotic use(18, 19). The decreased risk associated with male gender is unexplained; but independent associations for ESBL-EC/KP colonization have been described for both genders in previous studies(15, 20, 21).

Of particular importance was the high prevalence of chromosomal integration of *bla*_CTX-M_ in *E. coli* in this study (>14%), perhaps contributing to the stable propagation of this resistance gene family within certain strains. Chromosomal integration of *bla*_CTX-M_ in *K. pneumoniae* was not observed in our study, although it has been seen in Spain(22). In addition, despite the limitations of short-read assemblies, the genetic contexts of *bla*_CTX-M_ suggested high levels of genetic plasticity in flanking structures, and significant associations with IS*26* for *bla*_CTX-M-15_, *bla*_CTX-M-55_, and *bla*_CTX-M-27_. IS*26* has been previously hypothesised to facilitate the mobility of *bla*_CTX-M_ and genetic rearrangement of resistance gene plasmids, and is likely contributing to the dissemination of these resistance genes within the human gastrointestinal reservoir(23-25).

This study has several limitations. Our survey dates from 2012, and the epidemiology of ESC-R EC/KP carriage may have changed in the intervening timeframe; nevertheless, our data represent the largest molecular epidemiological study of gastrointestinal ESC-R-EC/KP colonisation in Cambodia and a useful benchmark for future studies. We only included up to three bacterial colonies per faecal sample, likely resulting in significant under-estimation of the diversity present at the population level(26). Short-read sequencing resulted in limited information regarding the wider genetic context of important resistance genes conferring ESC-R; nevertheless, we were still able to ascertain that the genetic contexts of these resistance genes are extremely diverse. Our outpatient study population may not be truly representative of healthy children in the community, given that these individuals had presented to the outpatient department for some form of medical review. Lack of more detailed information on some potential risk factors meant we were unable to fully assess the specific mechanisms promoting ESC-R EC/KP colonisation. Further work characterising the role of healthcare admissions, socio-economic factors and intestinal parasites on the acquisition and long-term carriage dynamics of these strains would be valuable. In addition, our sample size was too small and sparse to investigate geographical clustering of strain types, and to investigate specific risk factors for colonisation with common strain types or resistance gene alleles.

Despite these limitations, our study adds to the growing body of literature demonstrating widespread gastrointestinal colonisation with ESC-R-EC and ESC-R KP in Southeast Asia(8), and showing that exposure to this reservoir may in turn act as a source for the wider, global transfer of these strains(27). The genetic contexts of important resistance genes are highly mosaic, consistent with rapid exchange of resistance genes within and between bacterial hosts. Significant levels of chromosomal integration of the most important ESC-R gene family, *bla*_CTX-M_, were also observed, and may result in these genes being stably maintained and propagated in one of the most common community-associated pathogens, namely *E. coli*.

## MATERIALS AND METHODS

### Patients and setting

Faecal samples were obtained from a consecutive subset of children/adolescents (<16 years) who had been enrolled prospectively in a helminth prevalence study at Angkor Hospital for Children in Siem Reap, Cambodia, from 3^rd^ April 2012 to 29^th^ June 2012, as described in(28).

### Microbiological methods

Samples were frozen at -80°C as aliquots homogenised in 0.9% sterile saline with 10% glycerol within an hour of receipt in the laboratory. For this study, faecal samples were thawed, and aliquots diluted 1:10 in saline and incubated for 16 hours at 37°C on Orientation CHROMagar (BD, Oxford, United Kingdom) with 10 µg cefpodoxime and 10 µg imipenem discs (Oxoid, Basingstoke, United Kingdom). For each faecal sample, up to three pink and/or dark blue colonies with different colonial morphotypes that grew within the cefpodoxime zone of inhibition (presumed ESC-R-EC and ESC-R-KP respectively) were selected for further analysis. Each selected colony was tested using the British Society of Antimicrobial Chemotherapy (BSAC) combination disc method to identify whether cefpodoxime (ESC) resistance was mediated via ESBLs (Class A: cefpodoxime-resistant, and cefpodoxime+clavulanic acid-sensitive) or via non-ESBL mechanisms (e.g. Class C AmpC beta-lactamases: cefpodoxime-resistant, and cefpodoxime+clavulanic acid-resistant)(29). All identified ESC-R colonies were stored frozen at -80°C in nutrient broth with 10% glycerol.

### Whole genome sequencing and sequence data processing

DNA was extracted from sub-cultured ESC-R isolates using a commercial kit (Fujifilm Quickgene, Japan) with an additional mechanical lysis step (Fastprep MP Biomedicals, USA). All isolates were sequenced using the Illumina HiSeq 2500, generating 150bp paired-end reads. Sequence data have been deposited in GenBank (project accession: PRJNA391054).

To identify single nucleotide variants (SNVs) reads were mapped to species-appropriate reference genomes (*E. coli* CFT073 [GenBank: AE014075.1] and *K. pneumoniae* MGH78578 [GenBank: CP000647.1]), and variants called as described previously(30). Alignments of variable sites were padded to the length of the reference genome using bases with the same %GC content as that observed within each dataset. Bootstrapped, maximum-likelihood phylogenies were reconstructed for each species using RaxML version 7.7.6(31), using a generalised time-reversible model and four categories of rate heterogeneity (./RAxML-7.7.6/raxmlHPC-PTHREADS-SSE3 -f a -s <input_alignment.phy> -m GTRGAMMA -p 12345 -c 4 -x 12345 -# 100 -n <output_raxml_rapid_bootstrap>). Phylogenies have been deposited as projects in MicroReact to enable an interactive assessment of geographic distribution of genotypes (*E. coli*: https://microreact.org/project/By8bf5ajg; *K. pneumoniae*: https://microreact.org/project/Hy_yQcaog(32).

Contigs were assembled using Velvet/VelvetOptimiser (hash value range: 75-149)(33, 34). *In silico* MLST was determined by BLASTn(35) matches (100% match) to the Achtman/Pasteur MLST schemes for *E. coli* and *K. pneumoniae*(36, 37), and supported correct species identification. The presence/absence of resistance genes was determined using BLASTn and an in-house curated resistance gene database of over 60 gene families(38). Genes were considered present if a blast match of ≥80% of the query sequence was identified at ≥80% sequence identity using the *de novo* assemblies as blast databases. Ambler class genotype was class A if *bla*_CTX-M_, *bla*_SHV_ and/or *bla*_VEB_ were present, and/or class C if *bla*_CMY-2_, *bla*_DHA_ and *bla*_ACT-like_ genes were present. Where patient faecal samples yielded ≥2 strains, all resistance genes were treated as a single entity within the individual‘s profile.

The genetic context of *bla*_CTX-M_ was examined by extracting the contigs containing these genes, and annotating these using PROKKA(39), combined with BLASTn and manual annotation with reference to mobile genetic elements in the ISFinder database(40). Gene locations were characterised as “chromosomal” if other annotations on the contig were only found in chromosomal contexts in the top 20 BLASTn hits when the contig was compared with bacterial sequences available in GenBank (using default parameters); “plasmid” if the other annotations matched only plasmid sequences; or unknown if these conditions were not met e.g. the assembled contigs were too short to verify this.

### Epidemiological analyses

Information regarding putative risk factors for ESC-R EC/KP colonisation (collected on a standardised form) included details on: gender, age, hospitalisation status, residence in Siem Reap province versus elsewhere, water source (river, rain, well, bottled, piped, boiled), domestic animals (cats, dogs, birds), livestock (chickens, ducks, pigs, cows or water buffalo), toilet availability, malnutrition, co-morbidities, presence/absence of diarrhoea, presence/absence of parasites (assessed within (28)), soap usage for hand-washing and school attendance. No details regarding antibiotic consumption were ascertained within the study, but previous work locally has shown that individuals are often ill-informed about the nature of any medications used and that 32% of outpatient attendees have evidence of urinary antimicrobial activity(41).

### Statistical analyses

Independent risk factors for carriage were identified from a multivariable, stepwise, logistic regression model based on complete cases and initially including all factors (backwards elimination using exit p<0.1 to reduce over-fitting). A final multivariable logistic model was then fitted including all cases for which complete information was available for the retained risk factors. Statistical analyses were performed using STATA version 14 (StataCorp, College Station, USA).

### Ethical Approval

The study was approved by the Institutional Review Board (IRB), Angkor Hospital for Children, and the Oxford Tropical Research Ethics Committee (OXTREC 12–12). Caregivers of all included participants gave informed consent for their child to participate in the helminth survey, and for the samples to be used more widely in additional studies approved by the IRB.

## Acknowledgements

The authors wish to thank the staff and patients at Angkor Hospital for Children, Siem Reap, Cambodia, and members of the Modernising Medical Microbiology Informatics Group.

This work was supported by the National Institute for Health Research (NIHR) Oxford Biomedical Research Center (BRC). JvA is currently funded through a National Institute for Health Research (NIHR) Academic Clinical Fellowship. NS is currently funded through a PHE/NIHR/University of Oxford Clinical Lectureship; the sequencing work was also partly funded through a previous Wellcome Trust Doctoral Research Fellowship (#099423/Z/12/Z). TEAP and DWC are NIHR Senior Investigators.

The funders had no role in study design, data collection and interpretation, or the decision to submit the work for publication. The views expressed are those of the author(s) and not necessarily those of the NHS, the NIHR or the Department of Health.

The authors have no conflicts of interest to declare.

